# Paclitaxel causes *de novo* induction of invasive breast cancer cells by repolarizing tumor-associated macrophages

**DOI:** 10.1101/2025.01.13.632767

**Authors:** Madeline Friedman-DeLuca, George S. Karagiannis, Camille L. Duran, Robert J. Eddy, Suryansh Shukla, Jiufeng Li, John S. Condeelis, Maja H. Oktay, David Entenberg

**Affiliations:** Department of Pathology, Albert Einstein College of Medicine/ Montefiore Medical Center, Bronx, NY, USA; Integrated Imaging Program for Cancer Research, Albert Einstein College of Medicine/ Montefiore Medical Center, Bronx, NY, USA; Cancer Dormancy Institute, Albert Einstein College of Medicine/ Montefiore Medical Center, Bronx, NY, USA; Department of Microbiology and Immunology, Albert Einstein College of Medicine/ Montefiore Medical Center, Bronx, NY, USA; Gruss-Lipper Biophotonics Center, Albert Einstein College of Medicine/ Montefiore Medical Center, Bronx, NY, USA; Tumor Microenvironment and Metastasis Program, Albert Einstein College of Medicine/ Montefiore Medical Center, Bronx, NY, USA; Department of Surgery, Albert Einstein College of Medicine/ Montefiore Medical Center, Bronx, NY, USA; Department of Cell Biology, Albert Einstein College of Medicine/ Montefiore Medical Center, Bronx, NY, USA

**Keywords:** Breast cancer, macrophages, MenaINV, NF-κB, Notch1, paclitaxel, chemotherapy

## Abstract

Metastasis is the leading cause of breast cancer death, and tumor cells must migrate and invade to metastasize. Breast cancer cells that express the pro-metastatic actin regulatory protein MenaINV have an enhanced ability to migrate and intravasate within the primary tumor and extravasate at secondary sites. Though chemotherapy improves patient survival, treatment with paclitaxel leads to upregulation of MenaINV and an increase in metastasis in mice. MenaINV expression can be induced in breast cancer cells through cooperative NF-κB/ Notch1 signaling with macrophages, which are often increased in tumors in response to chemotherapy. MenaINV-expressing cells are also resistant to paclitaxel, raising the question of whether paclitaxel increases MenaINV by *de novo* induction or by selectively killing non-MenaINV-expressing cells. We hypothesized that paclitaxel causes *de novo* MenaINV induction by increasing macrophage-tumor cell NF-κB/ Notch1 signaling. Understanding this pro-metastatic effect of chemotherapy is crucial to refining treatment strategies.

In this study, we demonstrate that paclitaxel increases MenaINV expression by *de novo* induction. Mechanistically, paclitaxel induces MenaINV by repolarizing tumor-associated macrophages towards a pro-inflammatory phenotype. These pro-inflammatory macrophages then participate in enhanced NF-κB/ Notch1 signaling with tumor cells, which leads to MenaINV induction in the tumor cells. These results lay the groundwork for novel microenvironment-based therapies to alleviate the pro-metastatic effects of chemotherapy in breast cancer.

## Introduction

Breast cancer (BC) is the second leading cause of cancer death among women in the United States, with most deaths resulting from metastasis (1). To metastasize, tumor cells must migrate and intravasate within the primary tumor and extravasate at secondary sites. Tumor cells accomplish these tasks using invadopodia – actin-rich protrusions with proteolytic activity that degrade extracellular matrix. In BC cells, the actin regulatory protein MenaINV supports invadopodium stability and degradative activity (2), enhancing cell migration and metastasis. Furthermore, tumor cells that express MenaINV are more sensitive to chemotactic gradients in the tumor microenvironment, which facilitate invadopodium formation and directional migration towards blood vessels where they intravasate (3, 4). Indeed, BC cells that express MenaINV have an enhanced ability to migrate, intravasate, and extravasate (4-6), and high MenaINV expression is associated with poor outcomes in both mouse (4) and human (7, 8) BC.

Though chemotherapy improves patient survival, neoadjuvant paclitaxel increases the percent MenaINV+ area in both mouse and human residual breast tumors, as well as lung metastasis in mice (9). MenaINV-expressing cells are resistant to paclitaxel (10), so the observed increase in MenaINV after paclitaxel treatment may be due to the selective killing of non-MenaINV-expressing cells. However, we have shown that MenaINV is induced in tumor cells via cooperative NF-κB/Notch1 signaling with macrophages (11). Chemotherapy-induced tissue damage has been shown to cause a macrophage influx into tumors, potentially increasing macrophage-tumor cell NF-κB/Notch1 signaling and MenaINV induction. Thus, in addition to the selective killing of non-MenaINV-expressing cells, paclitaxel may also cause *de novo* induction of MenaINV. We hypothesized that paclitaxel induces MenaINV expression by increasing macrophage infiltration and macrophage-tumor cell NF-κB/Notch1 signaling. Here, we find that paclitaxel causes macrophages to adopt a pro-inflammatory phenotype that promotes NF-κB/Notch1 signaling and leads to *de novo* MenaINV induction in tumor cells.

## Results

### Paclitaxel causes *de novo* MenaINV induction

To test whether paclitaxel causes *de novo* MenaINV induction, we treated FVB-PyMT mice bearing spontaneous breast tumors with paclitaxel and co-administered either DAPT – to prevent MenaINV induction (11, 12) – or vehicle control (**Fig. 1A, Additional file 1: Materials and Methods**). DAPT is a γ-secretase inhibitor used to block Notch signaling. However, given the crosstalk between the Notch and NF-κB pathways, we (11) and others (13) have found that DAPT also inhibits NF-κB activation. If paclitaxel increases the percent MenaINV+ area only by selection, then dual DAPT/paclitaxel treatment would result in the same MenaINV increase as paclitaxel alone through the selective killing of non-MenaINV-expressing cells.

**Figure 1.**
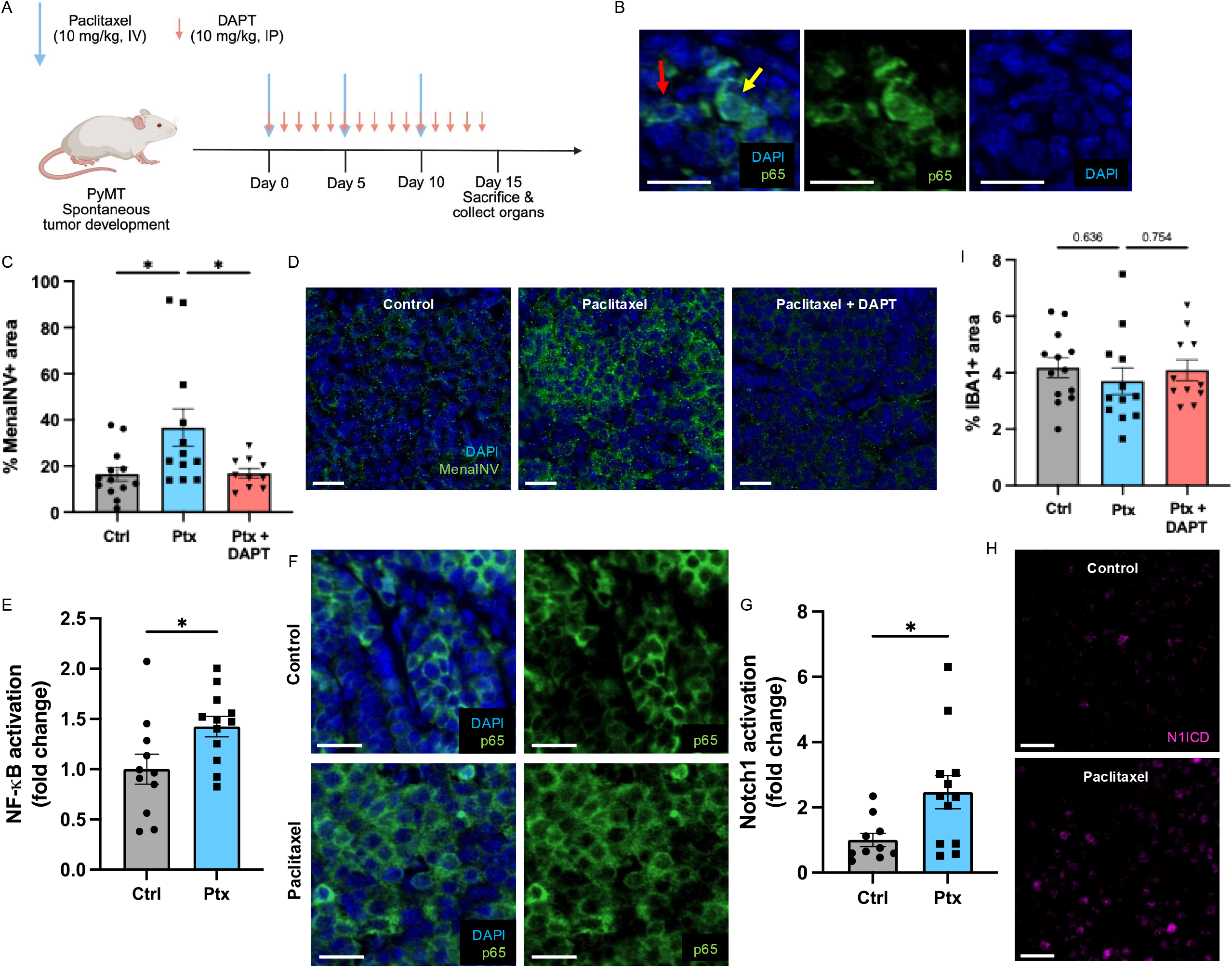
Paclitaxel increases MenaINV through *de novo* induction: (**A**) Experimental design of paclitaxel and DAPT treatments in PyMT mice. Figure created with BioRender.com. **(B)** Representative image of PyMT breast tumor stained for p65. Yellow arrow points to cell with active NF-κB signaling (nuclear p65). Red arrow points to cell with only inactive NF-κB signaling (cytoplasmic p65). (**C, D**) Quantification and representative images of MenaINV (measured by % MenaINV+ area) in tumors from mice treated with control (ctrl), paclitaxel (ptx), and paclitaxel + DAPT (ptx + DAPT). (**E, F**) Quantification and representative images of NF-κB activation (measured by p65 nuclear localization) in tumors from mice treated with control and paclitaxel. (**G, H**) Quantification and representative images of Notch1 activation (measured by expression of cleaved Notch1 intracellular domain) in tumors from mice treated with control and paclitaxel. **(I)** Macrophage infiltration (measured by % IBA1+ area) in tumors from mice treated with control, paclitaxel, and paclitaxel + DAPT. Data are reported as mean +/-SEM. Data in **C** and **I** were analyzed using a one-way ANOVA. Data in **E** and **G** were analyzed using a *t*-test. \**p* < 0.05. Scale bars = 20 μM. Each point on the graphs represents the average value from 10 fields of view taken from an individual mouse.

Following treatment, mice were sacrificed, and their tumors were analyzed for MenaINV expression, NF-κB activation (measured by p65 nuclear localization (**Fig. 1B**)), Notch1 activation (measured by cleaved Notch1 intracellular domain (N1ICD)), and macrophage infiltration. As previously observed, paclitaxel-treated tumors expressed significantly more MenaINV than controls (**Fig. 1C, D**, first two bars/panels), and this was accompanied by a significant increase in NF-κB and Notch1 activation (**Fig. 1E-H**). When MenaINV induction was inhibited with DAPT, paclitaxel no longer increased MenaINV (**Fig. 1C, D**, third bar/panel). This indicates that paclitaxel causes *de novo* MenaINV induction.

We originally hypothesized that this induction resulted from increased macrophage infiltration following paclitaxel treatment, but no difference was observed (**Fig. 1I**). This led us to question the necessity of macrophages in paclitaxel-mediated MenaINV induction.

### Macrophages are required for paclitaxel-mediated MenaINV induction

To test the role of macrophages in paclitaxel-mediated MenaINV induction, we used two mouse models of BC: FVB-PyMT mice as above, and SCID mice bearing the triple-negative HT17 patient-derived xenograft (14). Mice received paclitaxel and either clodronate liposomes – to deplete macrophages – or PBS control liposomes (**Fig. 2A; Additional file 2: Fig. S1A**). After treatment, tumors were analyzed for MenaINV expression, NF-κB and Notch1 activation, and macrophage infiltration. Paclitaxel again caused a significant induction of MenaINV, and this induction was prevented when macrophages were depleted (**Fig. 2B-D; Additional file 2: Fig. S1B, C**). This indicates that macrophages *are* required for paclitaxel to induce MenaINV. NF-κB and Notch1 signaling were also significantly decreased when macrophages were depleted during paclitaxel treatment (**Fig. 2E-H; Additional file 2: Fig. S1D, E**), indicating that macrophages are the primary source of additional NF-κB and Notch1 stimulation in the tumor microenvironment following paclitaxel treatment. Our data indicate that for paclitaxel to induce MenaINV, macrophages must be present, but not necessarily in higher proportions than in untreated conditions. This led us to ask whether paclitaxel changes the phenotype of macrophages to promote NF-κB/Notch1 signaling.

**Figure 2.**
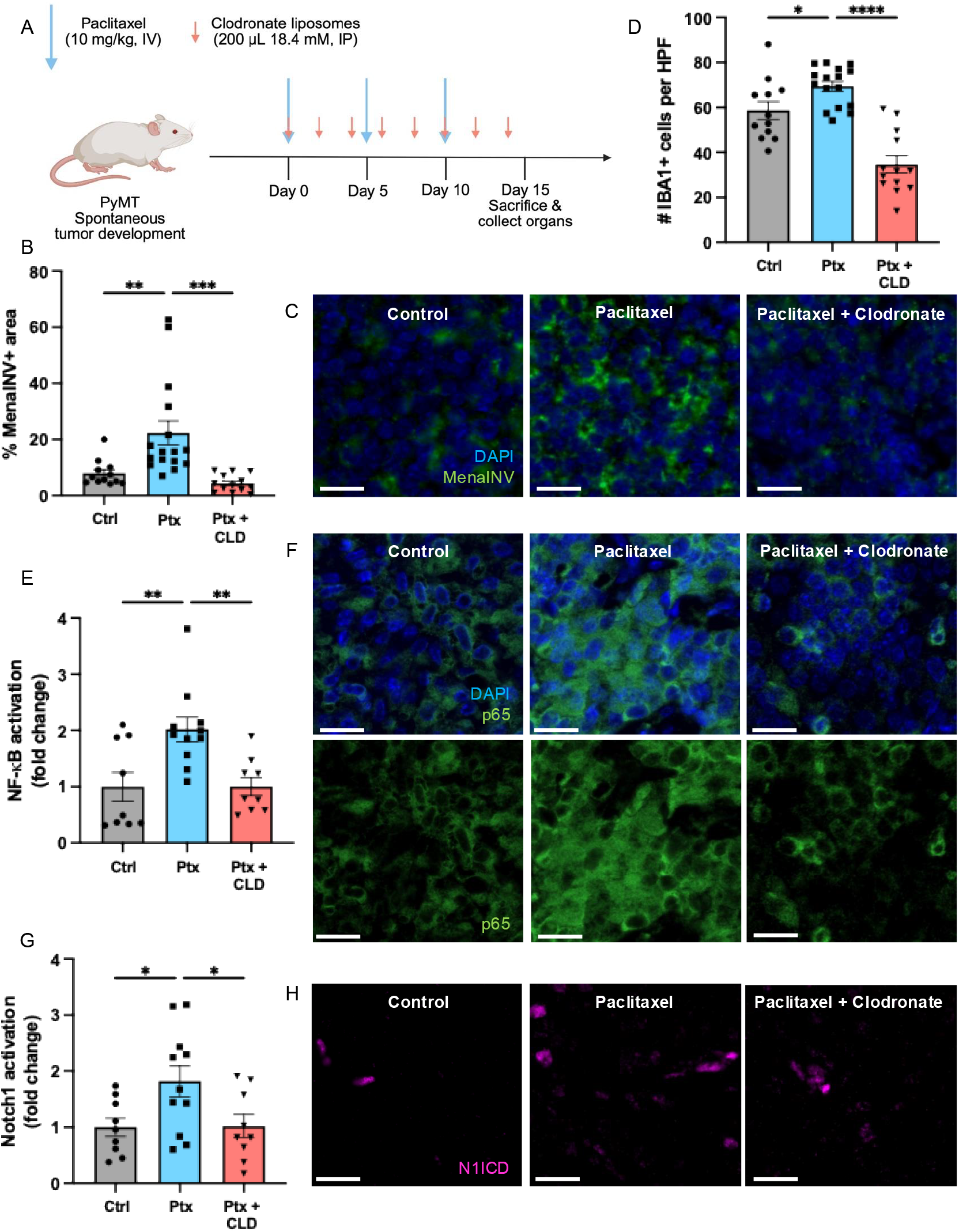
Macrophages are required for paclitaxel-mediated MenaINV induction: (**A**) Experimental design of paclitaxel and clodronate treatments in PyMT mice. Figure created with BioRender.com. (**B, C**) Quantification and representative images of MenaINV in tumors from mice treated with control (ctrl), paclitaxel (ptx), and paclitaxel + clodronate liposomes (ptx + CLD). (**D**) Macrophage infiltration (# IBA1+ cells per high power field (HPF)) in tumors treated with control, paclitaxel, and paclitaxel + clodronate. (**E, F**) Quantification and representative images of NF-κB activation in tumors from mice treated with control, paclitaxel, and paclitaxel + clodronate liposomes. (**G, H**) Quantification and representative images of Notch1 activation in tumors from mice treated with control, paclitaxel, and paclitaxel + clodronate liposomes. Data are reported as mean +/-SEM. Data were analyzed using a one-way ANOVA. \**p* < 0.05, \*\**p* < 0.01, \*\*\**p* < 0.001, \*\*\*\**p* < 0.0001. Scale bars = 20 μM. Each point on the graphs represents the average value from 10 fields of view taken from an individual mouse.

### Paclitaxel induces a pro-inflammatory phenotype in macrophages, which promotes NF-κB/Notch1 signaling

To investigate how paclitaxel affects the ability of macrophages to participate in NF-κB/Notch1 signaling, we cultured RAW264.7 murine macrophages or THP-1 human macrophages with 10 μM paclitaxel for 24 h and measured mRNA expression of Notch ligand Jagged1 and secretion of TNFα, IL-1β, and TGF-β1, which activate NF-κB. TGF-β1 can also enhance Notch signaling in several contexts (15). Paclitaxel significantly increased Jagged1 expression and TNFα/IL-1β secretion in both cell lines (**Fig. 3A-C; Additional file 2: Fig. S2A-C**), and TGF-β1 secretion in RAW264.7 cells (**Fig. 3D**), but not THP-1 cells (**Additional file 2: Fig. S2D**). These results indicate that paclitaxel causes macrophages to adopt a pro-inflammatory phenotype that promotes their participation in NF-κB and Notch1 signaling, though the exact phenotypic changes may differ between species.

**Figure 3.**
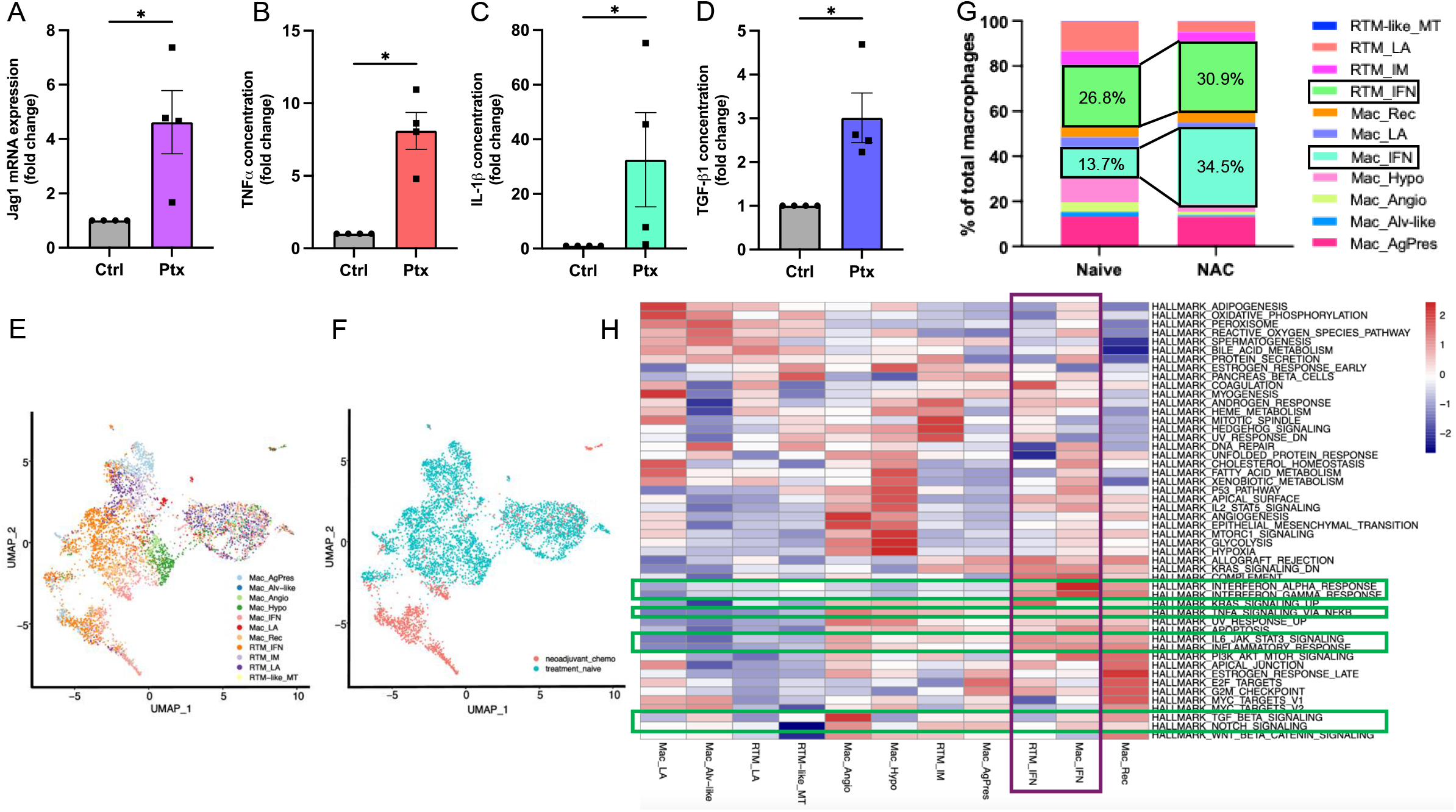
Paclitaxel alters macrophage phenotype, promoting NF-κB and Notch1 signaling: (**A**) Jagged1 mRNA expression in RAW264.7 macrophages treated with 10 μM paclitaxel. Secretion of (**B**) TNFIZl, (**C**) IL-1β, and (**D**) TGF-β1 from RAW264.7 macrophages treated with paclitaxel. (**E-H**) single-cell RNA-sequencing analysis of macrophages in the tumors of breast cancer patients who were treatment-naïve (naïve) or had undergone neoadjuvant chemotherapy (NAC). (**E**) UMAP (uniform manifold approximation and projection) of tumor-associated macrophages colored by macrophage subtype. Subtypes beginning with “Mac” are monocyte-derived macrophages (based on expression of monocyte markers *CCR2, VCAN, S100A6*, and *CD52*). Subtypes beginning with “RTM” are tissue-resident macrophages (based on expression of tissue residency markers *TCF12, MS4A4A, GATM, LYVE1, FOLR2, and PLTP*, as described in the original publication (17)). AgPres = antigen presenting, Alv-like = alveolar-like, Angio = angiogenic, Hypo = hypoxia-related, IFN = interferon-primed, LA = lipid-associated, Rec = recruited, IM = interstitial macrophage, MT = metallothioneins. (**F**) UMAP of tumor-associated macrophages colored by treatment group. (**G**) Stacked bar graph showing each macrophage subtype as a percentage of all macrophages in tumors of treatment-naïve and NAC-treated BC patients. NAC results in increases in the Mac_IFN and RTM_IFN subtypes (black outlines). (**H**) Gene set variation analysis (GSVA) shows that the macrophage subtypes that dominate the NAC group (Mac_IFN and RTM_IFN, magenta outline) have increased activation of pro-inflammatory pathways involved in NF-κB and Notch signaling (green outlines). Data in **A-D** are reported as mean +/-SEM and were analyzed using the Mann-Whitney U test. \**p* < 0.05. Each replicate was normalized to its control. All *in vitro* experiments were repeated 4 times.

### Neoadjuvant chemotherapy alters macrophage subtype composition in patients

We next investigated whether chemotherapy also repolarizes macrophages in BC patients. We identified a previously published dataset of single-cell RNA sequencing reads from human BC patient biopsies taken either before treatment (treatment-naïve) or after neoadjuvant chemotherapy (NAC), as described in the original publication (16). We annotated the macrophage compartment of this dataset based on ontogeny and functional gene signatures, as previously described (17), allowing our analysis to account for a spectrum of macrophage polarizations (**Fig. 3E**). UMAP visualization showed clear clustering between treatment groups, indicating distinct transcriptional profiles (**Fig. 3F**).

Both treatment groups exhibited macrophage heterogeneity. However, the macrophage population in NAC tumors was dominated by interferon-primed monocyte-derived (Mac_IFN) and tissue-resident (RTM_IFN) macrophages, which are characterized by high expression of interferon-primed genes, including *CCL8, IDO1*, and *CXCL9* (17) (**Fig. 3G**). The Mac_IFN subtype increased from 13.7% of macrophages in the treatment-naïve group to 34.5% of macrophages in the NAC group, while the RTM_IFN subtype rose from 26.8% to 30.9%. Together, these interferon-primed macrophages account for 65.4% of all macrophages in NAC tumors, compared to 40.5% in treatment naïve tumors.

Gene set variation analysis (GSVA) revealed that these two subtypes were highly upregulated in pathways related to inflammatory cytokine secretion (**Fig. 3H**). The Mac_IFN subtype was also upregulated in Notch signaling genes, which include the Notch ligands (**Fig. 3H**). These results confirm that NAC promotes an inflammatory phenotype in macrophages in human BC, which increases their ability to participate in NF-κB/Notch1 signaling, and complement our previous finding that NAC increases MenaINV expression in human BC (9).

Taken together, our results reveal that chemotherapy skews macrophages towards a pro-inflammatory phenotype, increasing their ability to participate in NF-κB/Notch1 signaling with tumor cells, which subsequently leads to MenaINV induction (**Fig. 4**).

**Figure 4.**
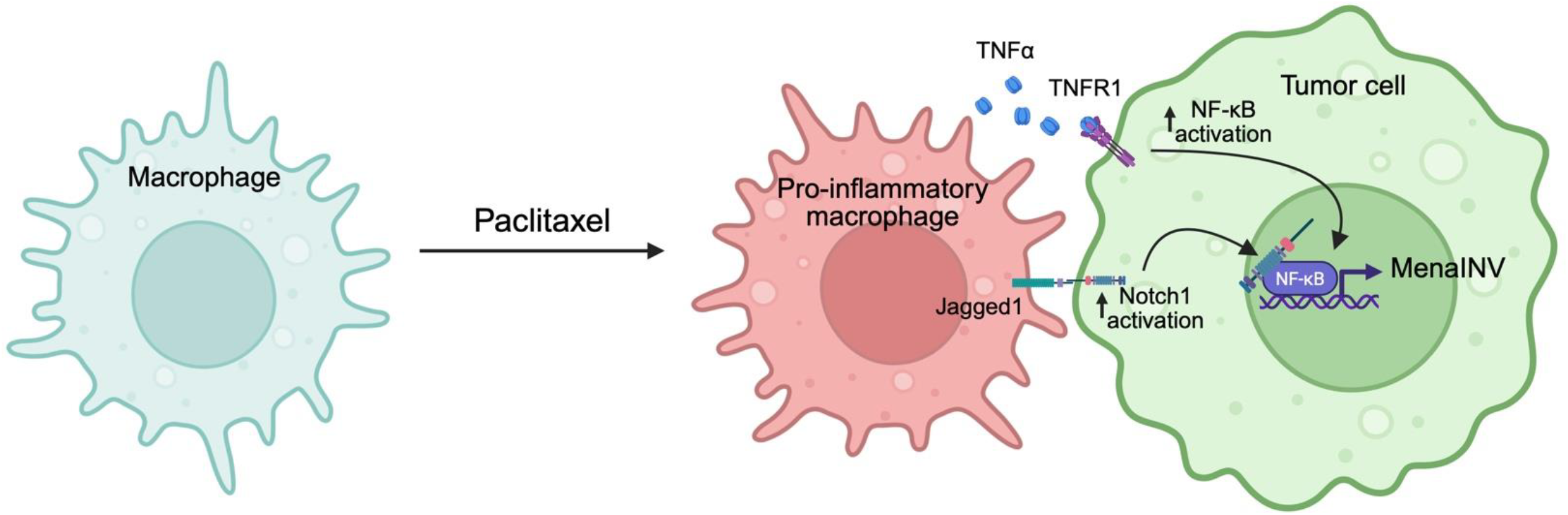
Paclitaxel skews macrophages towards pro-inflammatory phenotype, increasing macrophage-tumor cell NF-κB/Notch1 signaling, which induces MenaINV. Paclitaxel causes macrophages to take on a pro-inflammatory phenotype. Pro-inflammatory macrophages express higher levels of TNFIZl and Jagged1, which activate NF-κB and Notch1 signaling in tumor cells, respectively. Increased levels of NF-κB and Notch1 activation in tumor cells leads to increased MenaINV expression. Figure created with BioRender.com.

## Discussion

In this study, we discovered that neoadjuvant paclitaxel induces the expression of the pro-metastatic protein MenaINV in BC cells by promoting NF-κB/Notch1 signaling with macrophages. We previously observed a paclitaxel-driven increase in MenaINV (9, 11). However, because MenaINV expression confers resistance to paclitaxel (10), it was unknown whether this increase resulted from *de novo* induction or selective killing of non-MenaINV-expressing cells. Here, we show that the paclitaxel-driven increase in MenaINV is the result of *de novo* induction; when the pathways involved in MenaINV induction were blocked during paclitaxel treatment, no increase in MenaINV was observed (**Fig. 1C**). These results indicate that paclitaxel *induces* MenaINV expression.

We have previously shown that Notch1 signaling between macrophages and tumor cells is crucial for robust MenaINV induction (11, 12). The Notch1 receptor and its ligands are membrane-bound, so Notch1 signaling only occurs between adjacent cells. Given this and our previous finding that macrophages are necessary for the paclitaxel-mediated MenaINV increase (11), we hypothesized that paclitaxel causes a macrophage influx into the tumor, increasing opportunities for macrophage-tumor cell Notch1 signaling. Interestingly, we found that, while macrophages are necessary for paclitaxel to induce MenaINV (**Fig. 2B-D**), an increase in macrophage proportion is not (**Fig. 1C, I**). We conclude that, rather than increasing macrophage-tumor cell NF-κB/Notch1 signaling by increasing macrophage infiltration, paclitaxel repolarizes macrophages, allowing them to participate in such signaling more readily (**Fig. 3**). This aligns with previous reports that paclitaxel activates TLR4 receptors on macrophages, increasing inflammatory signaling (18), and complements our previous finding that TNFα increases MenaINV expression in tumor cells *in vitro* (11).

Chemotherapy has long been known to exert both pro- and anti-tumoral effects on macrophages. Indeed, prior evidence links chemotherapy-induced pro-inflammatory macrophages with elevated innate and adaptive anti-tumoral immune response (19). Here, we show that chemotherapy-induced pro-inflammatory macrophages also have the pro-tumoral effect of inducing the pro-metastatic protein MenaINV in tumor cells. As drug discovery platforms and protein-targeting capabilities advance, it is crucial to investigate the exact mechanisms behind both the pro- and anti-tumoral responses to chemotherapy to refine treatment strategies.

## List of abbreviations

BC: breast cancer
N1ICD: Notch1 intracellular domain
NAC: neoadjuvant chemotherapy
Mac_IFN: interferon-primed monocyte-derived macrophages
RTM_IFN: interferon-primed tissue-resident macrophages
GSVA: gene set variation analysis.

## Supplementary Information

Additional file 1: Materials and Methods (.pdf)

Additional file 2: Supplemental figures 1-2 and supplemental figure legends (.pdf)

## Declarations

### Ethics approval and consent to participate

All procedures were conducted in accordance with National Institutes of Health regulations and approved by the Albert Einstein College of Medicine Institutional Animal Care and Use Committee.

### Consent for publication

Not applicable

### Availability of data and materials

The datasets used and/or analyzed during the current study are available from the corresponding author on reasonable request.

### Authors’ contributions

MFD, MHO, and DE performed conceptualization. MFD, GSK, CLD, SS, JL, and RJE provided methodology. RJE developed and validated the MenaINV antibody. MFD and JL carried out formal analysis. SS & DE provided software and computational image analysis. MFD, GSK, and CLD performed investigation. MFD wrote the manuscript. GSK, JSC, MHO, and DE performed funding acquisition. MHO and DE supervised the study. All authors read and approved the final manuscript.

## Acknowledgments

We thank the Histopathology Core Facility for their assistance with sample processing and the Analytical Imaging Facility for their imaging services. We thank Luis Rivera Sanchez for assistance with macrophage quantification and Ved Sharma for contributing to sample generation. We thank Junya Zhang for performing the scRNA-seq analysis and Dianne Cox for advising on macrophage biology and methodology.

