## Supplementary material for "Paclitaxel causes *de novo* induction of invasive breast cancer cells by repolarizing tumor-associated macrophages": Materials and Methods

### Materials & Methods

#### Animal models

All procedures were performed in accordance with National Institutes of Health regulations and approved by the Albert Einstein College of Medicine Institutional Animal Care and Use Committee. Transgenic mice expressing the polyoma virus middle-T (PyMT) antigen under the control of the mouse mammary tumor virus long terminal repeat (MMTV-LTR) (1) were bred in house and have palpable mammary tumors at approximately 6 weeks of age. The HT17 patient-derived xenograft (PDX) was transplanted into SCID mice as previously described (2). Only female mice were used in experiments.

#### *In vivo* treatments

For MMTV-PyMT mice, treatments began at 8-10 weeks of age. For SCID mice bearing HT17 PDXs, treatments began 6 weeks after tumor transplantation, when mice were 12 weeks of age.

Paclitaxel (Sigma) was reconstituted to a stock concentration of 10 mg/mL in 1:1 ethanol/Cremophor-EL (Millipore Sigma 238470) and stored in aliquots at -20 °C. Immediately before treatments, the stock solution was further diluted in PBS and given at a dose of 10 mg/kg intravenously (total injected volume 200 μL) for a total of 3 doses 5 days apart. Mice in the vehicle control groups received the same volume of ethanol/Cremophor-EL diluted in PBS on the same schedule.

DAPT (Sigma D5942) was reconstituted in 100% ethanol to a stock concentration of 20 mg/mL and stored in aliquots at -20 °C. Immediately before treatments, the stock solution was further diluted in corn oil and given at a dose of 10 mg/kg intraperitoneally (total injected volume 100 μL) daily for 14 days. Mice in the vehicle control groups received the same volume of ethanol in corn oil on the same schedule.

Clodronate liposomes (Encapsula Nano Sciences, SKU# CLD-8901) or PBS control liposomes were stored at 4 °C. Mice received 200 μL liposomes (concentration 18.4 mM) intraperitoneally every 2 days for 14 days.

Tissue fixing and staining

Following treatments, mice were sacrificed, and mammary tumors were removed and fixed in 10% formalin for 24-48 hours (tumor to formalin volume ratio of 1:7). Tissues were then paraffin-embedded and cut in 5 μm thick sections. Slides were deparaffinized by melting the paraffin at 60 °C for 1 h, followed by 2 x 10-min incubations in xylene. Slides were then rehydrated in ethanol and water, and antigen retrieval was performed in 1x citrate buffer (pH 6.0, Vector H3300) at 97 °C for 20 min in a steamer.

For cleaved Notch1 intracellular domain (cN1ICD) and p65 staining, slides were then incubated in 0.3% hydrogen peroxide in water for 10 min to quench endogenous peroxidase and then blocked in a buffer containing 1% bovine serum albumin, 1% normal goat serum, and 10% fetal bovine serum (FBS) in 0.05% PBS-Tween20 for 1 h at room temperature. Slides were then stained using the multiplex tyramide signal amplification (TSA) immunofluorescence assay (Akoya Biosciences) according to the manufacturer's instructions with the following primary antibodies: rabbit anti-cN1ICD (1:200, CST4147), rabbit anti-p65 (1:4000, CST8242). Briefly, slides were incubated in the appropriate primary antibody diluted in blocking buffer overnight at 4 °C. The next day, slides were washed 3 x 5 min in 0.5% PBS-Tween20 and then incubated in goat anti-rabbit-HRP diluted 1:2000 in blocking buffer for 1 h at room temperature. After washing 3 x 5 min with 0.5% PBS-Tween20, TSA was performed by incubating slides in Opal reagents (Akoya FP1487001KT and FP1496001KT) diluted 1:100 in amplification buffer for exactly 9 min. The process of antigen retrieval, blocking, antibody incubation, and TSA was repeated for each primary antibody. After the final wash, slides were incubated in DAPI for 5 min and mounted with ProLong Diamond Antifade Mountant (Invitrogen).

For MenaINV and IBA1 staining, following antigen retrieval, slides were incubated in blocking buffer (same as above) for 1 h at room temperature. The tissues were then incubated in the following primary antibodies diluted in blocking buffer overnight at 4 °C: chicken anti-MenaINV (0.25 μg/mL, generated by Covance as previously described (3)), rat anti-IBA1 (1:800, Invitrogen MA5-38266). The next day, slides were washed 3 x 5 min in 0.05% PBS-Tween20 and incubated in the following secondary antibodies diluted in blocking buffer for 1 h at room temperature: goat anti-chicken 555 (1:250, Invitrogen A32932), goat anti-rat 647 (1:1000, Invitrogen A21247). After washing 3 x 5 min in 0.05% PBS-Tween20, slides were incubated in DAPI for 5 minutes and mounted using ProLong Diamond Antifade Mountant (Invitrogen). All stained slides were imaged on the Pannoramic 250 Flash II digital whole slide scanner using a 20 x 0.75NA objective lens.

Image analysis

To analyze the % MenaINV+ area and % IBA1+ area, the digital whole slide scans of slides stained for MenaINV and IBA1 were uploaded into CaseViewer (3DHISTECH), and 10 different 40X fields of view representative of the entire tumor were chosen for each mouse from invasive, non-necrotic areas of the tumor. Tif files of each field of view were acquired and uploaded to FIJI (NIH). In FIJI, images were thresholded based on the negative staining control, and the area fraction (area of signal/total area of field) was calculated in each channel for each field of view. The average value of the 10 fields per mouse is reported.

To analyze Notch1 activation, the digital whole slide scans of slides stained for cN1ICD were uploaded into CaseViewer, and representative fields of view were chosen and uploaded to FIJI as described above. In FIJI, images were thresholded based on the negative staining control.

The area of cN1ICD signal was normalized to the total DAPI area in each field, and the average value of the 10 fields per mouse is reported.

To analyze NF-κB activation via p65 nuclear localization, the digital whole slide scans of slides stained for p65 were uploaded into CaseViewer, and representative fields of view were chosen and uploaded to FIJI as described above. In FIJI, images were thresholded based on the negative staining control, and two masks were made - one of the total p65 signal and one of the DAPI signal. p65 was considered nuclear if it overlapped with a DAPI-stained nucleus. Nuclear p65 was normalized to the total DAPI signal in the field, and the average value of the 10 fields per mouse is reported.

<u>*In vitro* treatments</u>

Paclitaxel used in the cell culture (ThermoFisher J62734.MC) was reconstituted in DMSO at a stock concentration of 0.01 M and stored in aliquots at -20 °C.

RAW264.7 cell line: In the morning, RAW264.7 cells were seeded at a density of 7.5 x $10^5$ cells/well in a 6-well plate in DMEM containing 10% FBS and 1% penicillin/streptomycin. That evening, the media was replaced with serum-starving DMEM (1% FBS, 1% penicillin/streptomycin). The cells were serum-starved overnight. The next morning, the media was replaced with fresh serum-starving media with the indicated concentrations of paclitaxel. Conditioned media from the cultured cells were collected after 24 h and stored at -80 °C until use.

THP-1 cell line: In the evening, THP-1 monocytes were seeded at a density of 8.0 x $10^5$ cells/well in a 6-well plate in RPMI containing 10% FBS and 1% penicillin/streptomycin (complete RPMI). To differentiate the monocytes into macrophages, 5 ng/mL PMA (phorbol 12-myristate-13-acetate) was added to the media for 48 h. The media was then replaced with complete RPMI without PMA, and the macrophages were rested in complete RPMI for 48 h. Cells were then serum-starved overnight in RPMI containing 1% FBS and 1% penicillin/streptomycin. The next morning, the media was replaced with fresh serum-starving RPMI with the indicated concentrations of paclitaxel. Conditioned media and lysates from the cultured cells were collected after 24 h and stored at -80 °C until use.

Enzyme-linked immunosorbent assay (ELISA)

Levels of TNF$\alpha$, IL-1$\beta$, and TGF-$\beta$1 in the conditioned media were determined using ELISA kits (R&D Systems, DY1679, DY410, DY401, DY210, DY201, DY240) according to the manufacturer's instructions. The concentrations of secreted proteins were interpolated from the standard curve measurements.

mRNA extraction and qRT-PCR

Total RNA was extracted from macrophages using the RNA Mini Plus Kit (Qiagen, #74134). Complementary DNA (cDNA) was generated from 1 μg of total RNA using the iScript cDNA Synthesis Kit (Bio-Rad, #1708891), according to the manufacturer's instructions. Quantitative reverse transcription PCR (qRT-PCR) was conducted using Power SYBR Green PCR Master Mix (Applied Biosystems, Thermo Fisher Scientific, #4367659) on a QuantStudio 3 real-time PCR system (Applied Biosystems, Thermo Fisher Scientific). Jagged1 expression levels were normalized to GAPDH. The following primers were used: mouse Jagged1 5'- TGCCTGCCGAACCCCTGTCATAAT -3', 5'- CCGATACCAGTTGTCTCCGTCCAC -3'; human Jagged1 5'- AATGGCTACCGGTGTGTCTG -3', 5'- CCCATGGTGATGCAAGGTCT -3'; mouse GAPDH 5'- ATCAAGAAGGTGGTGAAGCA -3', 5'- AGACAACCTGGTCCTCAGTGT -3'; human GAPDH 5'- ATCAAGAAGGTGGTGAAGCA -3', 5'- GTCGCTGTTGAAGTCAGAGGA -3'.

Single-cell RNA sequencing analysis

Single-cell RNA sequencing data was obtained from the publicly available site http://biokey.lambrechtslab.org. Raw sequencing reads can also be found in the European Genome-phenome Archive under study number EGAS00001004809 (4). Only cells taken from biopsies before anti-PD1 therapy (“Pre” condition) were considered in our analysis.

*Data processing and integration:* Sequencing count matrices and corresponding metadata from multiple breast cancer patient samples were imported into R. Analyses were performed using Seurat v5.1.0 (5). Cell-level metadata (e.g., patient ID, clinical annotations) were incorporated during Seurat object creation to facilitate downstream stratification. Individual Seurat objects originating from different count matrices were further merged to generate a single, integrated dataset for joint analysis. The merged Seurat object underwent standard preprocessing: log-normalization (NormalizeData), identification of highly variable genes (FindVariableFeatures), and scaling (ScaleData). During scaling, potential sources of technical and biological variability—including mitochondrial transcript proportion (percent.mito), ribosomal content (percent.ribo), total detected genes per cell (nFeature_RNA), and cell cycle scores (S.Score and G2M.Score)—were regressed out to mitigate confounding effects. Dimensionality reduction was performed with Principal Component Analysis (PCA) using the RunPCA function. The optimal number of principal components (PCs) used in analyses was inferred empirically by assessing the cumulative variance explained and inflection points in the PC spectra. Batch effect correction across samples was performed using the Harmony algorithm implemented via Seurat v5's IntegrateLayers function (6). The resulting Harmony-corrected embedding was utilized to identify nearest neighbors (FindNeighbors). Integrated clustering was performed employing the Leiden algorithm (resolution = 0.6) using the Harmony neighbor graph. Finally, Uniform Manifold Approximation and Projection (UMAP) embedding was computed based on the Harmony-corrected dimensions for visualization and further analyses.

*Macrophage subset annotation and analysis:* Cells annotated as "Myeloid_cell" originating from the "Pre" timepoint were subsetted from the integrated object. A previously published human tumor myeloid cell atlas served as an external reference (7). To annotate macrophage cell types, deep generative modeling via scANVI was employed with 50 training epochs, two 256-neuron hidden layers, and a latent dimension of 30 (8). This semi-supervised approach integrated atlas labels while modeling novel populations, yielding refined probabilistic cell type assignments. Cells labeled as macrophages or other myeloid subsets with low counts (<10 per type) were excluded for stability. These annotated macrophage subsets were then re-processed using SCTransform (regressing out technical covariates and cell cycle scores), PCA, UMAP (based on top 30 PCs), nearest neighbor graph construction (FindNeighbors), and clustering using the Louvain algorithm (FindClusters, algorithm=4, resolution=0.4). Finally, cell type composition across experimental cohorts was visualized using stacked bar plots showing percentages calculated per cohort.

*Gene set variation analysis:* Functional pathway analysis was conducted on the annotated macrophage subset (based on ScanVI predictions). Gene Set Variation Analysis (GSVA) was performed using the GSVA package (9). Gene expression data (data layer) was used as input, along with the Hallmark gene sets (H) from the Molecular Signatures Database (10). Mean GSVA scores were calculated for each identified cell cluster (or predicted cell type for macrophages). Results were visualized as heatmaps of Z-scaled mean GSVA scores per cluster/cell types, using R package pheatmap.
