## Supplemental figures 1-2 and supplemental figure legends for "Paclitaxel causes *de novo* induction of invasive breast cancer cells by repolarizing tumor-associated macrophages"

### Supplemental figure 1.

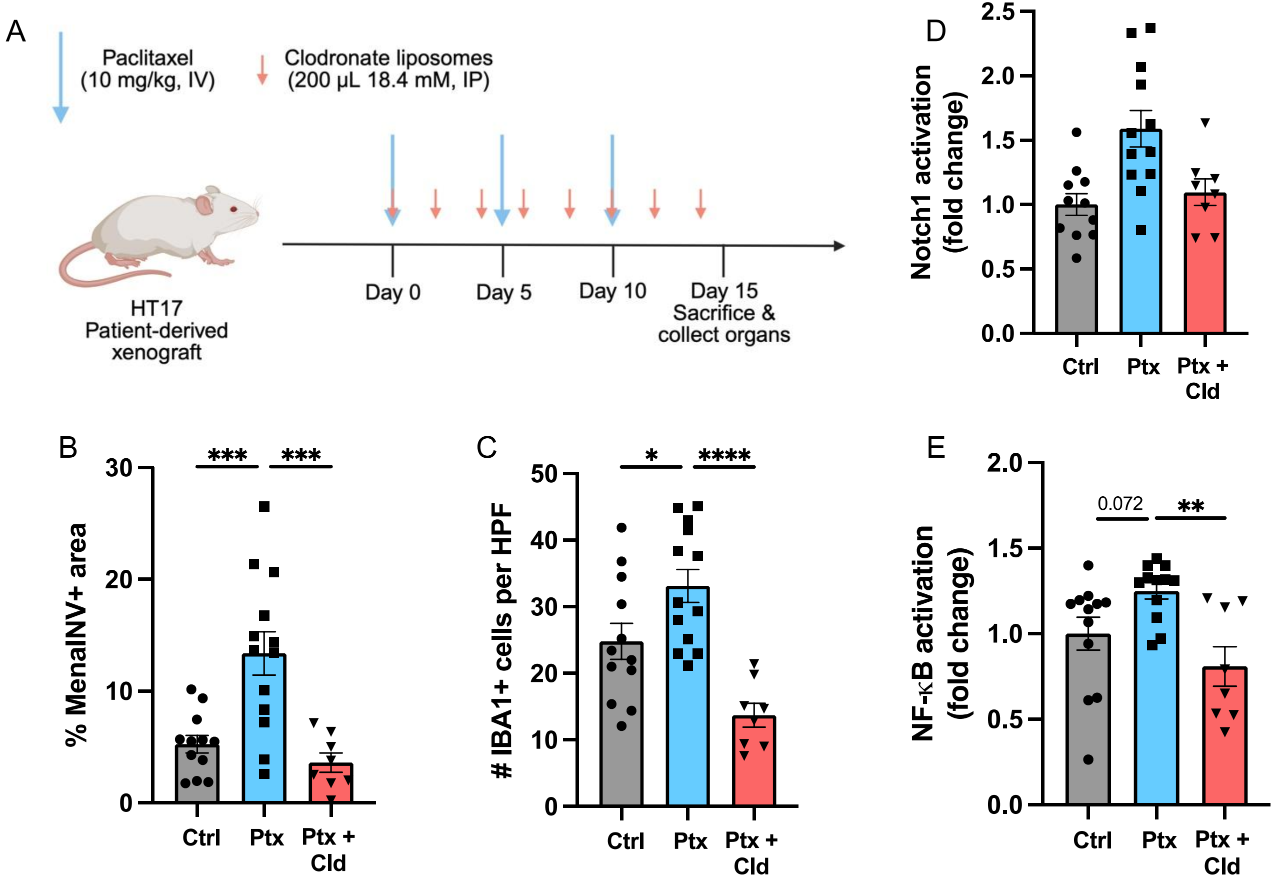

### Supplemental figure 2.

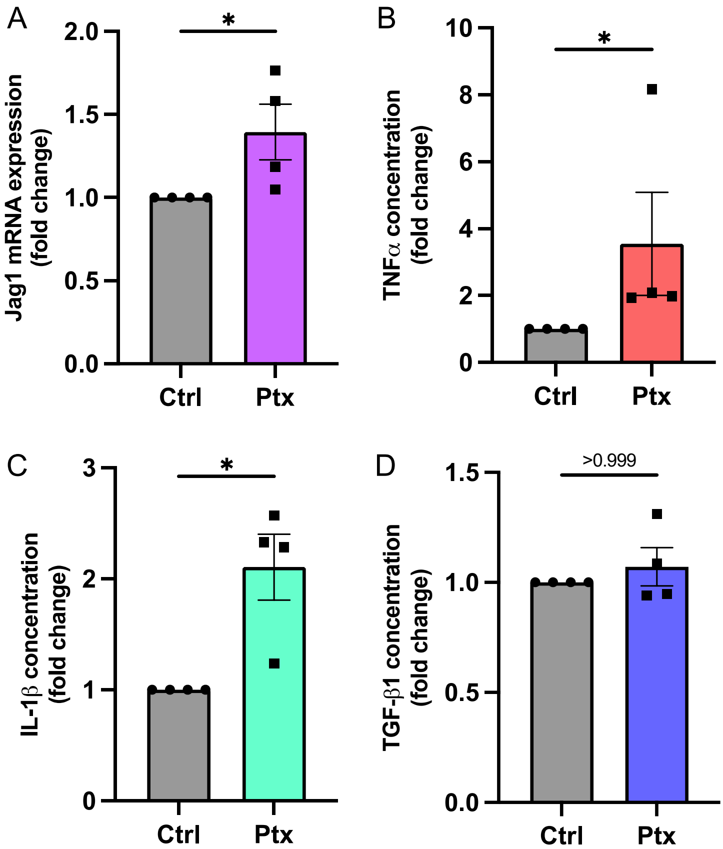

**Supplemental Figure Legends**

**Supplemental Figure 1. Macrophages are required for paclitaxel-mediated MenaINV induction in HT17 patient-derived xenograft model**: (**A**) Experimental design of paclitaxel and clodronate treatments in SCID mice implanted with the HT17 patient-derived xenograft. Figure created with BioRender.com. (**B**) % MenaINV+ area in tumors treated with control, paclitaxel, and paclitaxel + clodronate (cld). (**C**) Macrophage infiltration (# IBA1+ cells per high power field (HPF)) in tumors treated with control, paclitaxel, and paclitaxel + clodronate. (**D**) Notch1 activation in tumors treated with control, paclitaxel, and paclitaxel + clodronate. (**E**) NF-κB activation in tumors treated with control, paclitaxel, and paclitaxel + clodronate. Data are reported as mean +/- SEM. Data in **B-E** were analyzed using a one-way ANOVA. *$p < 0.05$, **$p < 0.01$, ***$p < 0.001$, ****$p < 0.0001$. Each point on the graphs represents the average value from 10 fields of view taken from an individual mouse.

**Supplemental Figure 2. Paclitaxel alters macrophage phenotype in vitro, promoting NF-κB and Notch1 signaling**: (**A**) Jagged1 mRNA expression in THP-1 macrophages treated with 10 μM paclitaxel. Secretion of (**B**) TNFα, (**C**) IL-1β, and (**D**) TGF-β1 from THP-1 macrophages treated with paclitaxel. Data are reported as mean +/- SEM and were analyzed using the Mann-Whitney U test. *$p < 0.05$. Each replicate was normalized to its control. All experiments were repeated 4 times.
